# Modular tissue-specific regulation of *doublesex* underpins sexually dimorphic development in *Drosophila*

**DOI:** 10.1101/585158

**Authors:** Gavin R. Rice, Olga Barmina, David Luecke, Kevin Hu, Michelle Arbeitman, Artyom Kopp

## Abstract

The ability of a single genome to produce distinct and often dramatically different male and female forms is one of the wonders of animal development. In most animals, sex-specific phenotypes are shaped by interactions between a sex determination pathway and spatial patterning gene networks. In *Drosophila melanogaster*, most sexually dimorphic traits are controlled by sex-specific isoforms of the *doublesex* (*dsx*) transcription factor, and *dsx* expression is mostly limited to cells that give rise to sexually dimorphic traits. However, it is unknown how this mosaic of “sex-naïve” and “sex-aware” tissues arises. Here, we characterize the *cis*-regulatory sequences that control *dsx* expression in the foreleg, which contains multiple types of sex-specific sensory organs. We find that separate modular enhancers are responsible for *dsx* expression in each sexually dimorphic organ. Expression of *dsx* in the sex comb is co-regulated by two enhancers with distinct spatial and temporal specificities that are separated by a genitalia-specific enhancer. Thus, the mosaic of sexually dimorphic and monomorphic organs depends on modular regulation of *dsx* transcription by dedicated cell type-specific enhancers.

**Summary Statement:** We identify the modular *cis*-regulatory elements that direct expression of *doublesex* in sexually dimorphic structures in *Drosophila* legs and genitalia. This regulatory landscape provides insight into how cells obtain their sex-specific identity.

## Introduction

Most animals are mosaics of sexually dimorphic and monomorphic tissues. Despite the multitude of traits that distinguish males from females, many tissues and organs lack overt sexual dimorphism. To understand how this mosaic pattern is produced, the action of sex determination pathways needs to be understood at the cellular level.

Sexually dimorphic morphologies are specified by diverse molecular mechanisms in different animal phyla (Kopp, 2012; Matson and Zarkower, 2012). One of the best studied mechanisms is found in the fruit fly, *Drosophila melanogaster*, where sex-specific development of most somatic cells is controlled by an alternative pre-mRNA splicing pathway (reviewed in Christiansen et al., 2002; Cline, 1993; Erickson and Quintero, 2007; McKeown, 1992). In *Drosophila*, the number of X chromosomes sets off a cell-autonomous cascade of sex-specific splicing that leads to the production of functional Transformer (Tra) protein in females, but not in males. Tra controls alternative splicing of *doublesex* (*dsx*) pre-mRNAs, such that the presence of functional Tra in females leads to a female-specific *dsx* isoform, and the absence of functional Tra leads to the production of a male-specific isoform (*dsx*^*F*^ and *dsx*^*M*^, respectively). *dsx* encodes transcription factors involved in the establishment of almost all morphological traits that differ between males and females (Baker and Ridge, 1980; Hildreth, 1965). The Dsx^M^ and Dsx^F^ proteins share a common DNA binding domain and bind to the same target sequences, but have different and sometimes opposite effects on gene expression, leading to sex-specific cell differentiation (Arbeitman et al., 2004; Arbeitman et al., 2016; Burtis et al., 1991; Goldman and Arbeitman, 2007; Lebo et al., 2009; Li and Baker, 1998; Yang et al., 2008). For instance, the *yolk protein 1, bric-à-brac*, *Fmo-2*, and *desat-F* genes are directly regulated by Dsx and are expressed at higher levels in females compared to males*. dsx* mutants show intermediate expression of *yolk protein 1, bric-à-brac*, and *Fmo-2* in both sexes, indicating that Dsx^F^ is likely activating, and Dsx^M^ repressing, these genes (Coschigano and Wensink, 1993; Luo and Baker, 2015; Williams et al., 2008). *desat-F*, on the other hand, is activated by Dsx^F^ but is not affected by Dsx^M^ (Shirangi et al., 2009). At the morphological level, loss-of-function *dsx* mutants develop as intersexes with a mixture of male, female, and intermediate traits (Baker and Ridge, 1980; Hildreth, 1965).

The details of the *tra*/*dsx* splicing cascade were characterized in the 1980s (Boggs et al., 1987; Burtis and Baker, 1989; Butler et al., 2018; McKeown et al., 1987; McKeown et al., 1988; Nagoshi et al., 1988), and are a textbook example of the role of alternative splicing in development. In contrast, the role of transcriptional regulation of *dsx* in *Drosophila* sexual differentiation was realized only recently. Although it may seem that every cell would need to “know” its sex, *dsx* is only transcribed in cells that are associated with sexually dimorphic organs (Camara et al., 2008; Rideout et al., 2010; Robinett et al., 2010; Tanaka et al., 2011). Thus, both males and females are mosaics of sexually dimorphic and monomorphic cells. This mosaicism makes it all the more remarkable that *dsx* regulates the development of so many different sex-specific traits, from pigmentation and genital morphology to brain neural circuits and gene expression in the gut (Goldman and Arbeitman, 2007; Keisman et al., 2001; Luo and Baker, 2015; Sanders and Arbeitman, 2008; Williams et al., 2008). To understand how this mosaic pattern is specified, we need to elucidate the mechanisms that establish *dsx* transcription.

Among the best studied sexually dimorphic organs in *Drosophila* are the foreleg bristles. The foreleg of *D. melanogaster* displays two distinct sex-specific features. First, the sex comb – a row of strongly modified mechanosensory bristles – is present only in males, while in females the homologous bristles retain the typical mechanosensory bristle morphology (Kopp, 2011; Tokunaga, 1962). Second, males have a greater number of foreleg chemosensory bristles compared to females (Mellert et al., 2012; Tokunaga, 1962). The sex comb and chemosensory bristles are found in close proximity to each other, and both are involved in mating behavior (Fan et al., 2013; Hurtado-Gonzales et al., 2015; Ng and Kopp, 2008; Spieth, 1952). The male-specific chemosensory bristles are used early in the stereotypic courtship sequence to taste the female cuticular pheromones (Fan et al., 2013; Spieth, 1952), while the sex comb assists in grasping the female prior to copulation (Hurtado-Gonzales et al., 2015; Ng and Kopp, 2008; Spieth, 1952).

*dsx* is required both for the distinctive sex comb morphology and for the higher number of chemosensory bristles in males (Belote and Baker, 1982; Mellert et al., 2012). Expression of *dsx* in the developing foreleg starts in the late 3^rd^ instar larva (Tanaka et al., 2011). During prepupal and early pupal development, this pattern resolves into several distinct clusters of cells corresponding to chemosensory bristle precursors, sex comb bristle precursors, and surrounding epithelial cells (Mellert et al., 2012; Robinett et al., 2010; Tanaka et al., 2011). This raises a question – is *dsx* expression in the foreleg controlled by a different, modular enhancer for each sex-specific bristle type, or by a common leg enhancer? More generally, is *dsx*, despite its apparently global function as a binary sex switch, subject to modular transcriptional control, with a dedicated enhancer for every cell population that expresses *dsx*?

To address this question, we identified and characterized the enhancers that control *dsx* expression in the sex comb and the foreleg chemosensory bristles. We found that three separate enhancers are responsible for regulating *dsx* expression in the foreleg. One enhancer drives broad foreleg expression that encompasses both sex comb and chemosensory organ primordia, while the other two have mutually exclusive activities – one in the sex comb, and the other in sex-specific chemosensory bristles. The two enhancers that contribute to sex comb development are separated by a genitalia-specific enhancer that has no activity in the leg. This complex *cis*-regulatory architecture suggests that *dsx* transcription, like that of other developmentally regulated transcription factors, is controlled by modular tissue-specific enhancers that can have both overlapping and non-overlapping spatial activities.

## Results

### Three enhancers with distinct spatial and temporal activities contribute to complex dsx expression in the foreleg

To identify the enhancers that specify the complex expression pattern of *dsx* during foreleg development, we generated a series of pPTGAL GAL4 reporter lines covering the nearly 50 kb of the upstream and intronic regions of the *dsx* locus (brown boxes, Fig. S1). In parallel, we examined an overlapping series of pBPGUw GAL4 reporters generated using a different vector and integration method (Pfeiffer et al., 2008) (grey boxes, Fig. S1). These lines were crossed to UAS-GFP.nls and screened 5 hr and 24 hr after puparium formation (APF). Using both sets of reporters, we identified three DNA fragments that drove expression patterns consistent with known *dsx* expression in the foreleg (Rideout et al., 2010; Robinett et al., 2010; Tanaka et al., 2011). We identified a chemosensory enhancer in a ~4 kb fragment of the first *dsx* intron (Fig. 1, F,G and Sup Fig. 1B, blue text), a sex comb enhancer in a ~3 kb section of the second intron (Sup Fig. 1C, red and green text), and an early foreleg enhancer in a ~4 kb section of the second intron downstream from the sex comb enhancer (Sup Fig. 1C, purple and green text). In parallel, another research group had identified a genital enhancer in the overlap between the 40F03 and 42D04 reporters (Sup Fig. 1C) (John Yoder, University of Alabama, personal communication), both of which showed foreleg expression in our screen. To test whether the sex comb and early foreleg enhancers were separate, and to reduce the temporal lag between enhancer activation and reporter detection, we cloned smaller sub-fragments of the *dsx* second intron into a nuclear GFP expression vector. We found that both the sex comb and the early foreleg enhancers drove the same expression patterns in this assay as in the larger GAL4 reporter constructs (Fig. 1 B-E).

**Figure 1:**
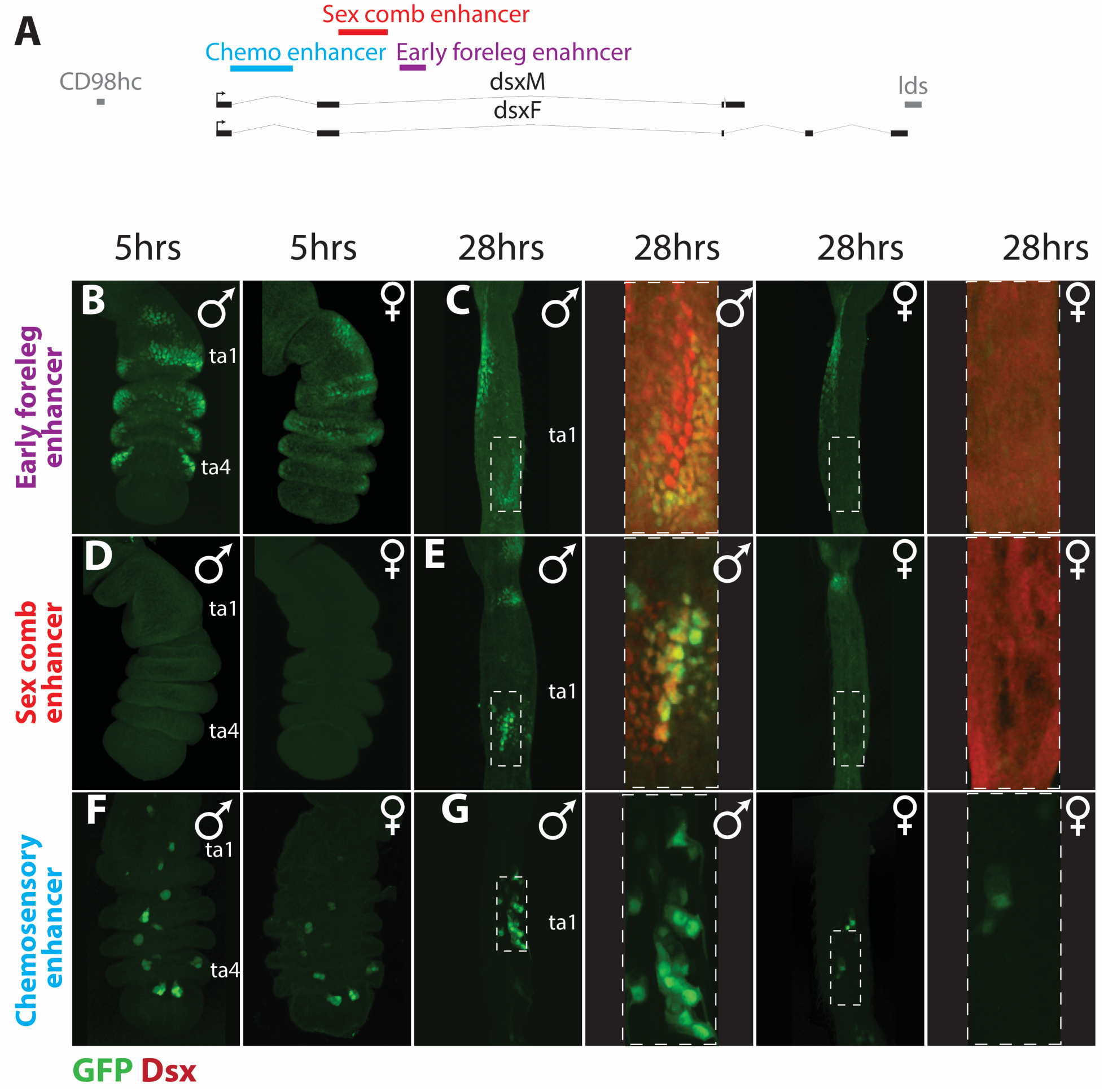
Three modular enhancers drive *doublesex* expression in the foreleg. **A)** A map of the *doublesex* locus with the positions of chemosensory (blue), sex comb (red), and foreleg (purple) *dsx* enhancers. Thick grey lines represent the flanking genes *lds* and *CD98hc*, thick black lines are the exons of *dsx*, and thin black lines are the *dsx* introns. Panels B – G show the foreleg expression patterns of the three *dsx* enhancers, with enhancer-driven GFP in green and magnifications of 28hr samples of the early foreleg and sex comb enhancers show Dsx antibody staining in red. Hours are after puparium formation (APF). The early foreleg and sex comb enhancers constructs are directly adjacent to GFP (Panels B-E), while the chemosensory enhancer (42C06 (Pfeiffer et al., 2008)) construct is a GAL4 driver crossed to a UAS-GFP.nls line. **B)** At 5 hours APF, the early foreleg enhancer recapitulates the Dsx expression pattern in the leg epithelium (Tanaka et al., 2011) and shows a similar expression pattern in males (left) and females (right). Both sexes show ectopic expression in the proximal first tarsal segment (ta1). **C)** At 28 hrs APF, the early foreleg enhancer shows clear sexual dimorphism in the first tarsal segment. Males show strong expression in the epithelial cells surrounding the sex comb and weak to no expression in the sex comb bristle cells. Females show little to no expression in the distal ta1. Both males and females show ectopic expression in the proximal ta1, in the same region as at 5 hrs APF (B), that is not seen by Dsx antibody staining (Tanaka et al., 2011). **D)** The sex comb enhancer is not active in either sex at 5 hrs APF, except for a proximal patch of ectopic expression seen near the joint in some individuals **E)** At 28 hrs APF, the sex comb enhancer is active in the bristle cells of the male sex comb (large nuclei), and weakly in the epithelial cells ventral to the sex comb. Females show weak expression in the distal portion of the segment. Both sexes show ectopic expression in the joint between the tibia and ta1. **F)** At 5 hrs APF, the chemosensory enhancer is active in small clusters of cells in ta1-ta5 in both sexes, with more GFP-positive cells in males than in females. **G)** At 28 hrs APF, both sexes show small clusters of expression in ta1-ta5; only ta1 is shown. (No Dsx antibody staining was performed in F-G).

The three enhancers direct distinct spatial and temporal expression of reporter genes, consistent with known *dsx* expression. *dsx* expression in the foreleg first becomes visible in wandering third instar larvae and is well developed in prepupal legs (Tanaka et al., 2011). Leg bristles are specified over several developmental periods. Most chemosensory bristles and the largest mechanosensory bristles are specified at the end of the 3^rd^ larval instar or early in prepupal development, while most mechanosensory bristles (including sex comb teeth) are specified between 6 and 15 hr APF (Belote and Baker, 1982; Held, 2002). At 5 hr APF, Dsx is present in an anterior-ventral region of tarsal segments 1 through 4 (ta1-4), where the sex comb and sex-specific chemosensory bristles will form (Tanaka et al., 2011). At this stage, the early foreleg enhancer drives reporter expression in a pattern similar to Dsx protein (Fig 1B). Although Dsx expression is already becoming sexually dimorphic by 5 hr APF (Tanaka et al., 2011), the GFP reporter intensity at this stage is roughly monomorphic, possibly due to the higher stability of the GFP protein (Fig 1B). The sex comb enhancer is not active at 5 hrs APF (Fig 1D), while the chemosensory enhancer is expressed in both sexes in small clusters of cells that match the pattern of developing chemosensory bristles (Fig 1F).

By 28 hr APF, Dsx expression is strongly dimorphic; in males, high Dsx expression is seen in sex comb bristles and surrounding epithelial cells, whereas in females, the homologous region of ta1 shows much lower Dsx expression (Tanaka et al., 2011; Fig 1C). In the rest of the tarsus, epithelial expression of Dsx disappears by 28 hr APF. At this stage in males, the early foreleg enhancer still shows weak activity in epithelial cells around the sex comb, especially on the distal-ventral side; however, it is not visibly active in sex comb bristles (Fig 1C). The sex comb enhancer drives a complementary pattern, with strong expression in sex comb bristles and weak expression in the surrounding epithelial cells (Fig 1E). In females, neither of these enhancers shows significant expression at 28 hr APF, consistent with the sexually dimorphic expression of Dsx (Fig 1C,E). For the chemosensory enhancer, cell clusters that reflect the pattern of chemosensory bristles continue to be observed at 28 hr APF in both males and females (Fig 1G). Thus, each of the three enhancers directs a distinct expression pattern in the foreleg.

### All three enhancers contribute to sex-specific sensory organ development

The patterns of GFP reporter expression suggest that the sex comb enhancer controls sex comb development and the chemosensory enhancer controls sex-specific chemosensory bristle development, while the early foreleg enhancer could potentially contribute to the development of both types of sex-specific bristles. To test these hypotheses, we used the chemosensory (42C06), foreleg (42D04), and sex comb (40F03) pBPGUw-GAL4 lines to drive a UAS-RNAi construct targeting the *shaven*/*Pax2* gene (*sv*). Knockdown of *sv* in sensory organ precursors causes severe truncation of bristle shafts (Kavaler et al., 1999), allowing us to identify the bristles where each *dsx* enhancer is active during prepupal and pupal development. We collected and analyzed 8 male and 5 female samples for each of the three GAL4 lines. This approach also allowed us to test for enhancer activity at developmental stages that were not observed by visual inspection of GFP reporters. The chaetotaxy of ta1 is particularly stereotypic and has been thoroughly mapped (Tokunaga, 1962), allowing us to compare the male and female knockdown phenotypes.

First, to identify all *dsx*-expressing bristles, we expressed UAS-*sv*RNAi using a *dsx*-GAL4 knock-in line, which was generated by inserting the GAL4 coding sequence into the *dsx* locus and reflects the full expression pattern of *dsx* (Robinett et al., 2010). We found that the ta1 chemosensory bristles fall into three classes (Fig 2). First, the dorsal-posterior chemosensory bristles are not affected by *dsx*-GAL4/UAS-*sv*RNAi; in the wild type these bristles are sexually monomorphic in size and position, suggesting they do not express or require *dsx* for their development (Fig 2A, B). Second, on the dorsal-anterior side, males have four chemosensory bristles on the distal half of ta1 (dark orange triangles in Fig 2A), compared to only one in females (Fig 2B); these bristles were always strongly affected by *sv* RNAi in both sexes, indicating that they express *dsx* (Fig 2A,B). The third class, two chemosensory bristles (yellow triangles in Fig 2A,B) were affected weakly and with more variability among individuals, suggesting that they may express *dsx* at lower levels or for a briefer period.

**Figure 2:**
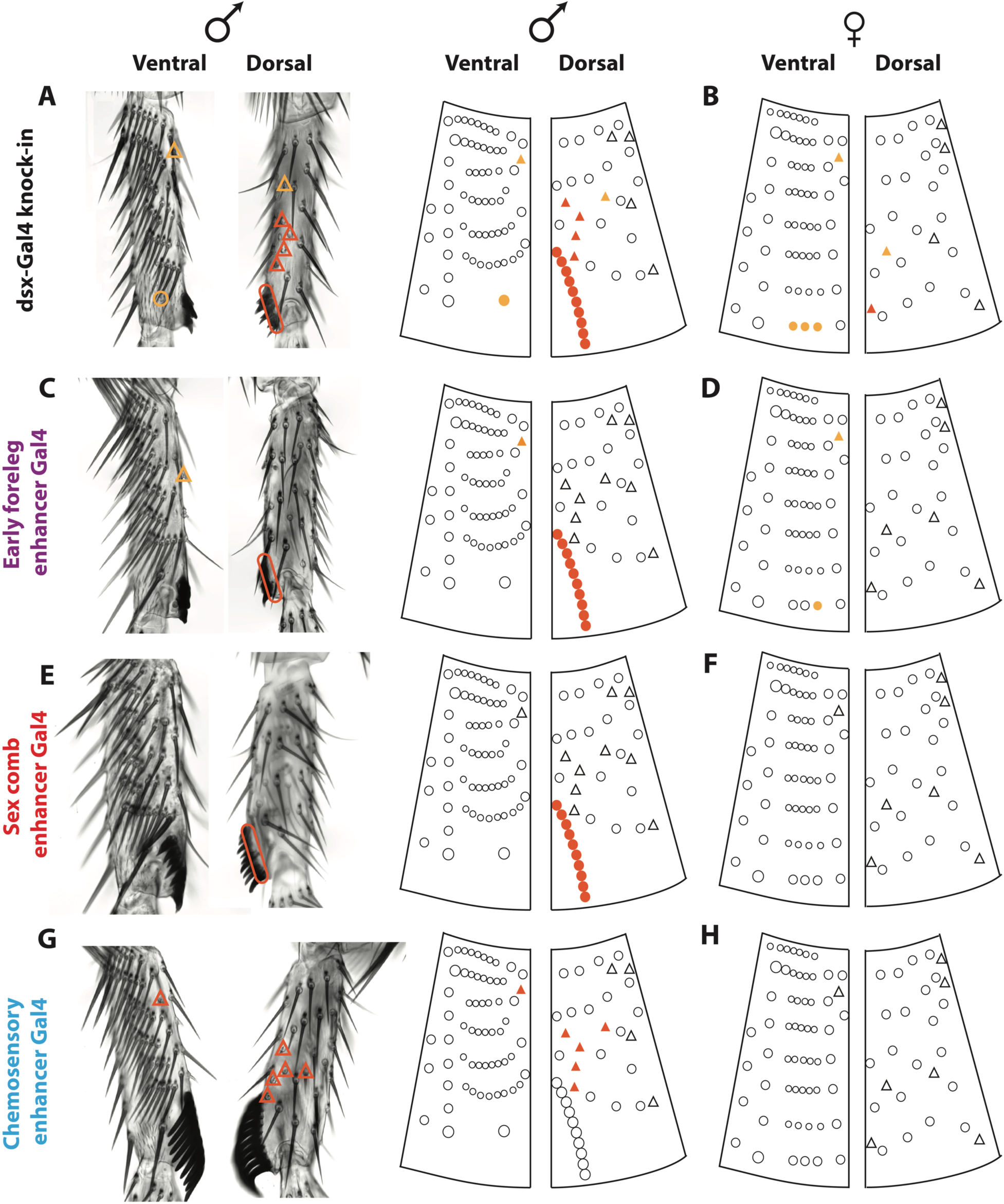
All three leg enhancers contribute to sex-specific bristle development. For each enhancer, a GAL4 reporter was used to drive a UAS-*shaven* RNAi construct, truncating the bristles where that enhancer is active. Images of male ta1 are shown next to a schematic representation of male and female ta1 chaetotaxy modified from (Tokunaga 1962). Circles designate mechanosensory bristles, while triangles designate chemosensory bristles. Dark orange symbols mark bristles that were affected in all individuals; bristles marked with yellow varied among individuals of the same genotype raised under standard conditions. **A)** A GAL4 knock-in in the *dsx* gene, which reflects the full expression pattern of *dsx* (Robinett et al., 2010), affects both sex comb and sexually dimorphic chemosensory bristles in males. **B)** In females, the *dsx*-GAL4 knock-in has a variable effect on the distal Transverse Bristle Row (TBR), which is homologous to the sex comb, and on some chemosensory bristles. **C)** In males, the early foreleg enhancer affects sex comb bristles in all flies as well as a single chemosensory bristle in some individuals. **D)** In females, the early foreleg enhancer affects one bristle of the distal TBR and the same chemosensory bristle as in males. **E)** In males, the sex comb enhancer affects only the sex comb. **F)** The sex comb enhancer does not affect any bristles in females. **G)** In males, the chemosensory enhancer affects the same chemosensory bristles as the *dsx*-GAL4 knock-in but has no effect on the sex comb. **H)** The chemosensory enhancer does not affect any bristles in females.

The male sex comb was always severely affected by *dsx*-GAL4/UAS-*sv*RNAi, whereas the homologous female bristles were affected weakly and with more variability (Fig 2A,B), consistent with a lower level of *dsx* expression in females compared to males (Tanaka et al., 2011). With the exception of the sex comb and homologous female bristles, no mechanosensory bristles were affected by *dsx*-GAL4/UAS-*sv*RNAi (Fig 2A,B), confirming their sexually monomorphic nature.

Next, we used each of the three *dsx* leg enhancers to drive UAS-*sv*RNAi and compared the resulting phenotypes in ta1 to that of the *dsx* knock-in. The early foreleg enhancer was similar to the *dsx* knock-in in having a strong and consistent effect on the male sex comb, but only a weak and variable effect on homologous bristles in females (Fig 2C,D). It also affected a single chemosensory bristle in proximal ta1 in both sexes (Fig 2C,D). As expected, the sex comb enhancer had a clear effect on sex comb development in males, but did not affect chemosensory bristles or any bristles in females (Fig 2E,F). The chemosensory enhancer consistently affected all anterior-ventral chemosensory bristles in males, including the two proximal bristles that were only weakly affected by the *dsx* knock-in; it did not affect any mechanosensory bristles (Fig 2G). Surprisingly, the chemosensory enhancer had no effect in females (Fig 2H), even though some female chemosensory bristles express *dsx* (Fig 2B), and the chemosensory enhancer drives UAS-GFP expression in ta1 in females (Fig 1C).

In addition to their function in ta1, the foreleg and chemosensory enhancers show expression in the more distal segments (ta2-ta4) (Fig 1B, F). These segments also contain chemosensory bristles, some of which are sexually dimorphic (Mellert et al., 2012; Tokunaga, 1962). In ta2 and ta3, the *dsx* knock-in and the chemosensory enhancer affect the same subset of four chemosensory bristles, while the sex comb enhancer has no effect (Fig Sup 2). The early foreleg enhancer affected some but not all of the bristles affected by the *dsx* knock-in, as well as one of the chemosensory bristles that was not affected by the *dsx* knock-in or the chemosensory enhancer (Fig Sup 2). Pupal expression of the early foreleg enhancer is epithelial, not bristle-specific (Fig 1E); thus, it is difficult to say whether this phenotype reflects a true developmental function of the early foreleg enhancer in sensory organs, or the perdurance of the GAL4 protein or the RNAi effect from an earlier time in development.

Examination of adult bristle phenotypes caused by UAS-*sv*RNAi expression suggests that the three leg enhancers of *dsx* make distinct contributions to sex-specific bristle development. The foreleg and sex comb enhancers function in sex comb development, while the chemosensory enhancer functions in the development of sexually dimorphic chemosensory organs. We also cannot rule out some contribution of the early foreleg enhancer to chemosensory bristle development.

### The foreleg and sex comb enhancers are modular and flank a separate genital enhancer

The foreleg and sex comb enhancers are located in close proximity within the second intron of *dsx* and are both active in the sex comb, although in different spatial patterns and at different times (Fig 1B-E). The same broad region also contains an enhancer that is active in the third instar larval genital disc (John Yoder, University of Alabama, personal communication). Mutations in *dsx* have been previously shown to disrupt clasper (also known as the surstylus) bristles (Hildreth, 1965), which have a morphology similar to sex comb teeth. Through antibody staining, we found that Dsx is expressed in several regions of the pupal male genitalia (Fig. S3). In the developing clasper, Dsx is enriched in the bristle precursor cells, similar to its enrichment in sex comb bristles. Analysis of the genital enhancer showed weak expression in the clasper as well as strong expression in the dorsal postgonites (Fig. S3, S4). These observations led us to inquire whether the foreleg, genital, and sex comb enhancers were fully modular and independent, or parts of a broader pleiotropic enhancer with overlapping activities in multiple tissues. To distinguish between these possibilities, we generated non-overlapping reporter constructs for each enhancer and examined each reporter both in the foreleg and in the genitalia. In addition, we generated larger constructs where either the sex comb or the early foreleg enhancer was excluded (Fig 3).

**Figure 3:**
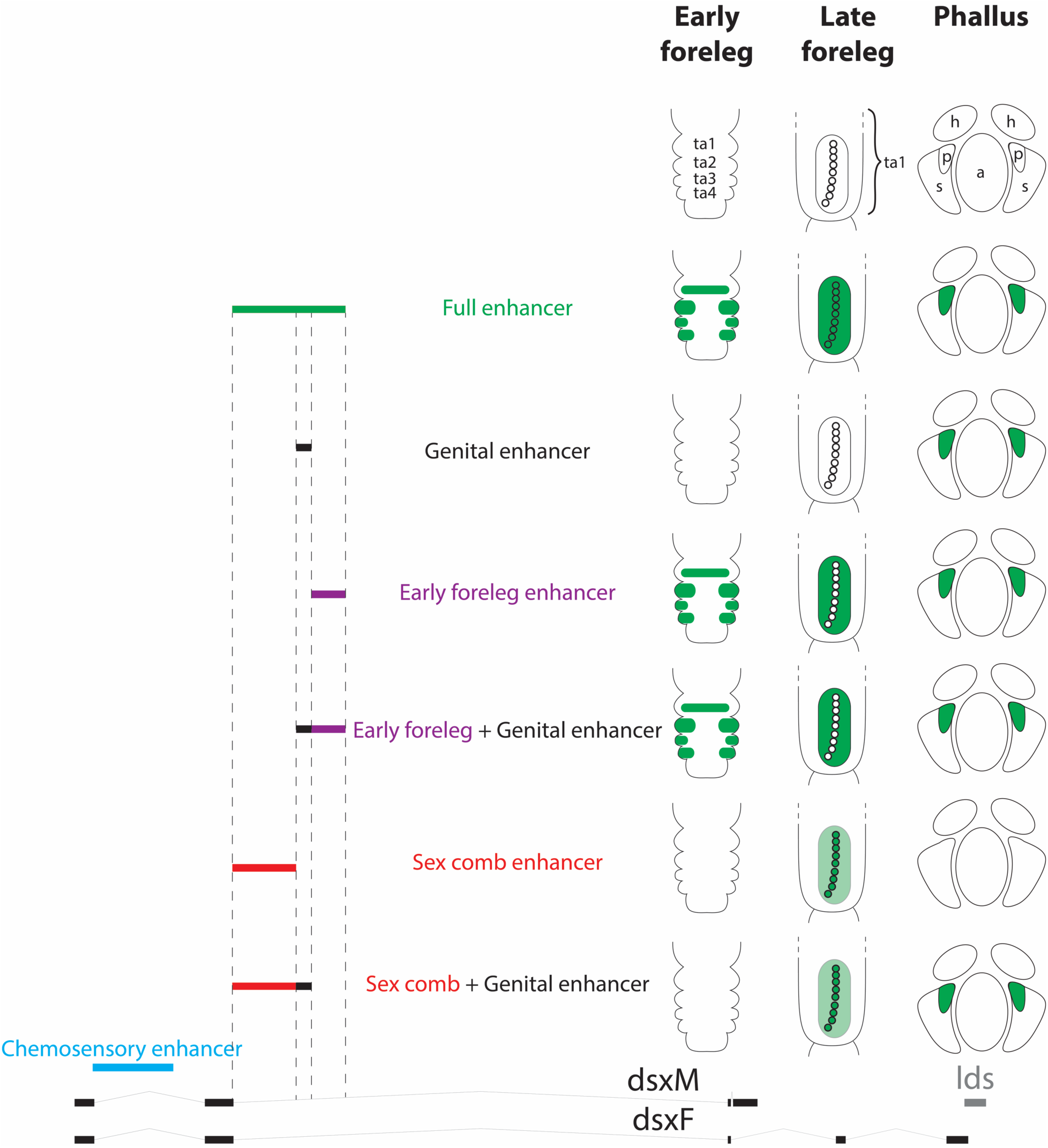
The foreleg and sex comb enhancers are separate and modular. To test for overlap between the foreleg, genital, and sex comb enhancer activities, the region they encompass was split into six constructs: full (all three enhancers, green), genital (black), foreleg (purple), foreleg + genital (purple + black), sex comb (red), and sex comb + genital (red + black). In the drawings, green shading shows reporter expression that matches endogenous Dsx expression, with light green indicating weak expression and white a lack of expression. Representative images of each construct are shown in Fig. S3. Only the constructs that include the early foreleg enhancer drive expression in the forelegs at 5 hrs APF and in the epithelial cells surrounding the sex comb at 24 hrs. Only regions that include the sex comb enhancer drive expression in the sex comb bristles at 24 hrs APF. Both the genital and the early foreleg enhancer constructs are able to drive expression in the genitalia at 48 hrs. ta1-ta4: tarsal segments 1-4; h: hypandrium; a: aedeagus; p: dorsal postgonites; s: aedeagal sheath.

We first confirmed that a 5.5 kb genomic fragment containing all three candidate enhancers (sex comb, genital, and foreleg) was able to drive both early and late expression in the foreleg, as well as genital expression (Fig. 3 and Fig. S4). Each separate enhancer can be seen as driving a distinct subset of that overall pattern. The late sex comb enhancer is active in the sex comb bristle cells but not in the genitalia, while the genital enhancer drives expression in the genitalia but has no activity in the foreleg (Fig. 3 and Fig. S4). The construct encompassing the early foreleg enhancer is active in the epithelial cells surrounding the sex comb, as well as in the genitalia in a pattern that resembles the genital enhancer (Fig. S4). However, further dissection of this region shows that the foreleg and genital activities are largely separable. The sequences necessary for accurate foreleg expression are located on the 3’ side of the foreleg enhancer and do not drive genital expression, while the sequences that activate expression in the genitalia are located on the 5’ side and only drive ectopic expression in the foreleg (Fig. 4 G-H and Fig. S3). Pairwise combinations of enhancers behave in a predictably modular fashion: a construct including the sex comb and genital enhancers drives genital and late sex comb expression, but no early epithelial expression in the foreleg, while the construct including the genital and early foreleg enhancers drives early foreleg and genital expression but no expression in sex comb teeth (Fig. 3 and Fig. S4). The chemosensory enhancer, located in a different intron and thus clearly distinct from the foreleg and sex comb enhancers (Fig 3). In summary, we conclude that the three leg enhancers are all fully separable, modular elements, but we cannot rule out some minor overlap between the foreleg and genital enhancers.

**Figure 4:**
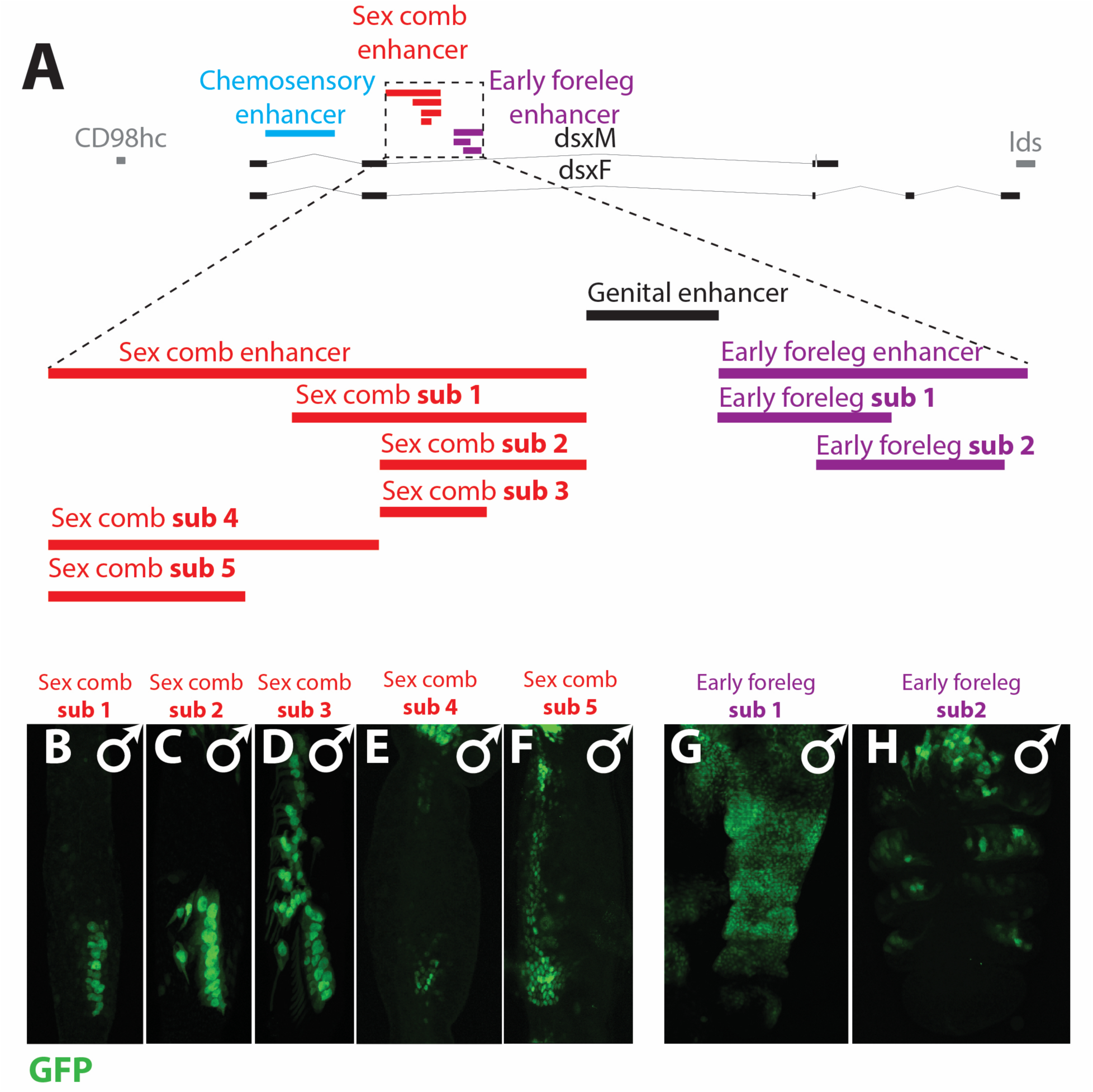
Repressor elements are necessary to limit *dsx* expression to sexually dimorphic cells. **A)** A map of the sex comb (red), genital (black), and foreleg (purple) enhancers. **B-E)** Representative images of male forelegs at 36 hrs APF for the *sex comb sub1-sub5* constructs; only the ta1 segment is shown. **B)** The *sex comb sub1* enhancer drives expression in the sex comb bristle cells; **C)** *sex comb sub2* is active in sex comb bristles but has additional ectopic expression in the central bristle and in the second most distal TBR; **D)** *sex comb sub3* is expressed in all TBRs bristle cells in the ta1 segment; **E)** *sex comb sub4* is expressed in some of the epithelial cells that surround the sex comb, but these cells are not directly adjacent to the sex comb bristles; **F)** *sex comb sub5* at 24 hrs AFP is expressed in the epithelial cells surrounding the sex comb, but also shows ectopic expression in more proximal epithelial cells **G-H)** Foreleg expression of the *foreleg sub1* and *sub2* constructs at 5 hrs APF. The *foreleg sub1* region drives broad ectopic expression throughout the ta1-ta4; *foreleg sub2* shows expression in a pattern similar to the complete early foreleg enhancer (Fig. 1B).

### Sequences in the foreleg and sex comb enhancers confine their activity to sexually dimorphic primordia

To identify the minimal sequences required for correct activity, we subdivided the sex comb and early foreleg enhancers into smaller reporter constructs (Fig 4A). While the ~3kb sex comb enhancer drives specific expression in sex comb bristles and the epithelial cells distal to the sex comb (Fig 1E), shorter sequences are unable to recapitulate this pattern. The 3’ ~1.5 kb of the sex comb enhancer (*sex comb enhancer sub1*) drives specific expression in sex comb bristles but is not expressed in the epithelial cells (Fig 4B). A shorter ~1 kb 3’ fragment (*sex comb enhancer sub2*) retains expression in the sex comb bristles, but also causes ectopic expression in the most distal TBR (Fig 4C). A core sequence of ~0.5 kb (*sex comb enhancer sub3*) expands expression to all ta1 TBRs (Fig 4D). The TBRs are serially homologous to the sex comb (Kopp, 2011; Tokunaga, 1962), but are sexually monomorphic and do not express Dsx (Robinett et al., 2010; Tanaka et al., 2011).

The 5’ ~1.5kb of the sex comb enhancer (*sex comb enhancer sub4*) drives expression in some of the epithelial cells surrounding the sex comb, but this expression does not encompass all of the epithelial cells where the full sex comb enhancer is active (Fig. 4E). Reduction of this sequence to a ~1kb 5’ fragment (*sex comb enhancer sub5*) causes ectopic expression in proximal epithelial cells (Fig 4F) that do not express Dsx (Robinett et al., 2010; Tanaka et al., 2011). Thus, sequences within the sex comb enhancer are necessary to prevent its activity in sexually monomorphic bristle and epithelial cells.

Dissection of the early foreleg enhancer identified a shorter 3’ fragment (*early foreleg enhancer sub2*) capable of driving expression in the correct region of the foreleg (Fig. 4H). A partially overlapping fragment (*early foreleg enhancer sub1*) shows broad ectopic activity throughout the tarsal segments (Fig 4G). Thus, similar to the late sex comb enhancer, both activating and repressive sequences are needed to limit this enhancer’s activity to sexually dimorphic cell populations.

## Discussion

### dsx expression in the foreleg is controlled by modular enhancers with both distinct and overlapping activities

In this study, we have characterized the *cis-*regulatory architecture directing the precise spatiotemporal and cell type-specific expression of *dsx* in the foreleg. Our results help explain the origin of the developmental mosaicism that generates a mix of sex-specific and sexually monomorphic organs in *Drosophila*, and possibly in other insects. Given the largely cell-autonomous control of insect sexual differentiation, the essential function of *dsx* enhancers is to provide spatial landmarks that establish which organs will become sexually dimorphic. In principle, *dsx* expression in different types of foreleg bristles could be controlled either by a single, broadly acting leg enhancer, or by multiple enhancers, each dedicated to a different bristle type. For example, the proneural genes *achaete* and *scute*, which control bristle specification, have very complex *cis*-regulatory regions with dozens of modular enhancers responsible for different, stereotypically patterned bristles (Garcia-Bellido and de Celis, 2009). Other enhancers are pleiotropic and active in multiple tissues, reflecting a history of evolutionary co-option from the more ancient organs to the more recently evolved (Glassford et al., 2015; Nagy et al., 2018; Rice and Rebeiz, 2019). In fact, some genes contain both modular (tissue-specific) and pleiotropic (multi-tissue) enhancers (Preger-Ben Noon et al., 2018).

We observe a partial overlap in the temporal and spatial activities of the three leg enhancers (Fig 1–3). However, these are not functionally redundant “shadow enhancers” of the type found in some other genes (Frankel et al., 2010; Hong et al., 2008). During prepupal development, the early foreleg enhancer is expressed in cells that give rise both to the sex comb and to the chemosensory bristles (Fig 2, Sup Fig 2). Later, the early foreleg enhancer expression in the sex comb becomes complementary to the activity of the sex comb enhancer: the former is expressed only in the epithelial cells, and the latter predominantly in sex comb bristles and only weakly in the epithelial cells. At pupal stages, when the early foreleg enhancer is no longer active in bristle cells, the late-acting sex comb and chemosensory enhancers have completely non-overlapping expression patterns. Overall, the existence of these three modular enhancers suggests that *dsx* expression in the foreleg goes through two successive stages: relatively broad expression early in development, followed by tightly limited, cell type specific expression later on.

These phases of *dsx* expression may have different developmental functions. For example, *dsx* knockdown in chemosensory bristles late in bristle development causes a defect in the axonal midline crossing by the bristle neurons (Mellert et al., 2012). Early *dsx* expression in the foreleg induces a male-specific increase in the number of bristles both in the sex comb and in the chemosensory system (Belote and Baker, 1982; Mellert et al., 2012). The broad early foreleg expression of *dsx* is established prior to the formation of mechanosensory bristles (Tanaka et al., 2011), suggesting that the early foreleg enhancer may control sex comb size by modulating the specification of sensory organ precursors. On the other hand, the later-acting sex comb enhancer is more likely to be involved in controlling the morphology of the individual sex comb teeth, which develop from mechanosensory bristle shafts but, in contrast to most such bristles, are thick, blunt, and darkly pigmented. Indeed, knockdown of *dsx* late in development affects the morphology but not the number of sex comb teeth, which become masculinized in females and feminized in males (Belote and Baker, 1982). Both the foreleg and the sex comb enhancers are active in the epithelial cells around the sex comb, suggesting that both could contribute to sex comb rotation – a coordinated sequence of cell shape changes that moves the sex comb from an initial transverse to the final longitudinal orientation (Atallah et al., 2009; Ho et al., 2018; Malagón et al., 2014; Tanaka et al., 2011). Since sex comb development involves so many different cellular processes spread over multiple developmental stages, it is perhaps not surprising that it requires multiple *dsx* enhancers.

Not all tissues and cell types behave in a modular fashion with respect to *dsx* expression. In the central nervous system, only one of the pPTGAL reporters tiling the *dsx* region showed an overlap with endogenous *dsx* expression (in 4 neurons in the PC2 cluster) (M. Arbeitman, unpublished). A separate study identified a *dsx* enhancer upstream of the transcription start with an activity overlapping Dsx protein expression in the brain (Zhou et al., 2014). However, most of the Dsx CNS expression pattern is not accounted for by any of the *dsx* reporter constructs (M. Arbeitman, unpublished). This suggests that for most *dsx*-expressing cells in the CNS, *dsx* regulation may not be due to modular enhancers acting independently, but rather to more complex, long-distance sequence interactions that are not captured by ~5 kb reporter fragments. If true, such “entangled” regulatory architecture could be more constraining than the modular enhancer organization we observe in the foreleg, which could in turn lead to slower evolution of *dsx* expression in the brain compared to other tissues.

### cis-regulatory information limits dsx expression to sexually dimorphic cells

In contrast to some enhancers where very short sequences are sufficient to convey correct regulatory information (Grieder et al., 1997; Manak et al., 1994; Swanson et al., 2010), both the sex comb and the early foreleg enhancers are quite large and cannot be reduced to a core region without perturbing their activities. Interestingly, dissecting these enhancers further leads not to the loss of activity, but rather to its expansion. This is similar to the abdominal enhancer of the *ebony* gene, where a core 0.7 kb sequence is sufficient for abdominal activity, but a much broader genomic region is necessary to prevent ectopic expression (Rebeiz et al., 2009). We find a similar pattern in our study: both the foreleg and the late sex comb enhancers of *dsx* have core sequences of less than 1 kb capable of driving expression in the foreleg, but require additional sequences to repress ectopic activity.

The late sex comb enhancer is particularly illuminating in this respect. The sex comb develops from the most distal transverse bristle row on ta1 (Atallah et al., 2009; Tanaka et al., 2009; Tokunaga, 1962), so the bristles that comprise it are serially homologous to the more proximal TBRs. However, while the distal TBR that gives rise to the sex comb undergoes dramatic male-specific modification, the remaining TBRs retain their original, sexually monomorphic morphology. Consistent with this, *dsx* is expressed, and the sex comb enhancer of *dsx* is active, only in the most distal TBR. Loss of sequences at the 5’ end of this enhancer leads to an expansion of its activity to the proximal, sexually monomorphic TBRs. Thus, much of the regulatory information contained in the sex comb enhancer is devoted to restricting its activity to sexually dimorphic cells, from a broader population of similar cells. This may reflect a general trend for *dsx* expression: in each tissue or body part (such as the brain, midgut, foreleg, body wall epidermis, etc.), only a subset of cells express *dsx* and undergo sex-specific differentiation (Goldman and Arbeitman, 2007; Luo and Baker, 2015; Rideout et al., 2010; Robinett et al., 2010; Sanders and Arbeitman, 2008; Tanaka et al., 2011). Integrating both activating and repressive inputs in *dsx* enhancers may be necessary to segregate these cells from otherwise similar but sexually monomorphic cells.

### Modular dsx enhancers may allow different sex-specific traits to evolve independently

While the sex combs are present in only a subset of *Drosophila* species, particularly in the *melanogaster* and *obscura* species groups, most though not all *Drosophila* species have sex-specific chemosensory bristles on their forelegs. Thus, one could imagine a more ancestral chemosensory foreleg enhancer becoming co-opted for sex comb development in the *melanogaster* and *obscura* groups. A more extreme scenario would involve a co-option of a genital enhancer for sex-specific foreleg development. Male genitalia and legs are serially homologous and share similar developmental programs involving some of the same transcription factors and signaling pathways (Gorfinkiel et al., 1999; Keisman et al., 2001; Sánchez et al., 1997), and the male genitalia of most *Drosophila* species have strongly modified clasper bristles reminiscent of sex comb teeth. Indeed, we find that Dsx is expressed in the clasper during late genital development. However, our results argue against a simple co-option scenario. The chemosensory bristle enhancer of *dsx* is clearly distinct from both enhancers that contribute to sex comb development, being located in a different intron. The situation is less clear with the early foreleg enhancer, where we cannot rule out a minor overlap with the genital enhancer. However, the late sex comb enhancer, which appears to play the dominant role in the development of sex comb teeth, is clearly separate from the genital enhancer.

Both the sex comb and the chemosensory bristles play important roles during courtship (Hurtado-Gonzales et al., 2015; Mellert et al., 2012; Ng and Kopp, 2008) and have undergone extensive changes in number, size, morphology, and spatial distribution in *Drosophila* evolution. The ability of all three foreleg enhancers to function and evolve independently of each other may have been a key element driving this evolutionary diversification. Analysis of these enhancers in other *Drosophila* species will provide important insights into the role of *dsx* in the evolution of sex combs and chemosensory bristles. The new GAL4 drivers that mark different types of sex-specific sensory organs may also prove valuable for dissecting the neural mechanisms of courtship behavior.

### Modular transcriptional regulation of dsx may reflect an evolutionarily ancient mode of sexual development

Dsx-related transcription factors (the *Dmrt* gene family) play important roles in sexual differentiation in animals as different as vertebrates, insects, and nematodes (Kopp, 2012; Matson and Zarkower, 2012). Outside of insects, however, *Dmrt* genes do not produce distinct male and female isoforms, but are regulated instead at the transcriptional level. Sex-specific splicing of *dsx*, and more generally the mode of sexual differentiation based on alternative splicing, is an insect innovation (Kopp, 2012; Wexler et al.). In crustaceans and chelicerates, *dsx* homologs are transcribed exclusively or predominantly in males, particularly in male-specific organs, and are required for male but not for female sexual development (Kato et al., 2011; Li et al., 2018; Pomerantz and Hoy, 2015). In *C. elegans*, the *Dmrt* genes *mab3, mab23*, and *dmd3* are expressed in tightly restricted cell lineages and are essential for male-specific differentiation of these cells, but are dispensable in females (Lints and Emmons, 2002; Mason et al., 2008; Ross et al., 2005; Serrano-Saiz et al., 2017; Yi et al., 2000). In vertebrates, the *Dmrt1* gene is highly expressed in the male but not in the female gonad, and is necessary to direct or maintain male-specific differentiation of the initially bipotential gonad (Matson et al., 2011).

These similarities suggest that sexual differentiation based on male-specific transcription of *Dmrt* genes in restricted cell populations is an evolutionarily ancient mode of establishing sexual dimorphism, and that in insects, sex-specific splicing of *dsx* was overlaid on this ancestral mechanism (Kopp, 2012; Wexler et al.). The finding that *dsx* transcription in *Drosophila* is controlled by multiple modular enhancers may reflect this more ancestral mode of sexual development. We predict that a similar modular control of *dsx* expression will be found in other arthropods, from basal insects and crustaceans to chelicerates.

## Materials and Methods

### Fly stocks

Fly strains were obtained from the Bloomington Stock Center, including UAS-GFP.nls (Bloomington stock #4775, genotype: w[1118]; P{w[+mC]=UAS-GFP.nls}14), UAS-svRNAi P{TRiP.JF02582}attP2 (Bloomington stock #27269, genotype: y[1] v[1]; P{y[+t7.7] v[+t1.8]=TRiP.JF02582}attP2), and *dsx*-GAL4 (Robinett et al., 2010) (Bloomington stock #66674, genotype: w[1118]; TI{GAL4}dsx[GAL4]/TM6B, Tb[1])

### Transgenic reporter constructs

Candidate enhancer regions were cloned into the GAL4 reporter vector pPTGAL (Sharma et al., 2002), and randomly integrated into the *D. melanogaster* genome by P-element transformation. Three independent lines from each of ten overlapping fragments were examined. In addition, we analyzed 15 independently generated GAL4 lines which span the same non-coding regions of *dsx* but have different fragment boundaries (Pfeiffer et al., 2008). These lines were constructed using the site-specific pBPGUw vector (Pfeiffer et al., 2008) and integrated into either the attP2 or attP40 genomic landing sites via the PhiC31 integrase system (Groth et al., 2004; Venken et al., 2006). Each line from both sets was crossed to a UAS-GFP.nls reporter and GFP expression was examined in the developing foreleg at 5 and at 24 hours after puparium formation (APF).

While the high level of GFP expression driven by the GAL4/UAS system was useful for the preliminary screening of genomic fragments, subsequent analysis was performed using direct GFP reporters, which enable more accurate temporal resolution. Genomic fragments were amplified by PCR, cloned into either pCR2 or pCR8 vectors (Invitrogen), transferred into the pGreenFriend GFP destination vector (Miller et al., 2014) using either restriction/ ligation or Gateway cloning, and integrated into the attP2 or attP40 genomic landing sites using the PhiC31 integrase system. Germline transformation was performed either in the Kopp lab or by BestGene (http://www.thebestgene.com/). In tests performed on a subset of constructs, we found little difference between attP2 and attP40; we subsequently used the attP40 site, which gave higher integration efficiency. In designing the sub-fragments of the sex comb and early foreleg enhancers, we used Evo-printer (Odenwald et al., 2005) and VISTA (Frazer et al., 2004) to identify sequences within these enhancers that were conserved between *D. melanogaster* and *D. pseudoobscura*. Whenever possible, we avoided breaking up highly conserved regions. The boundaries of all reporter fragments and the primers used to amplify them are listed in Supplementary Tables 1 and 2.

### Light Microscopy

Legs were dissected from adult flies and mounted in PVA-Mounting-Medium (BioQuip) or Hoyer’s media to clear the samples. Images were taken under Brightfield illumination using a Leica DM500B microscope with a Leica DC500 camera.

### Immunohistochemistry and Confocal Microscopy

Samples were aged, dissected, fixed, stained and imaged as described in Tanaka et al 2009 and Mellert, et al 2012. The primary mouse-Dsx[DBD] antibody (a gift from C. Robinett and B. Baker, later obtained from the Developmental Studies Hybridoma Bank, Cat# DsxDBD, RRID:AB_2617197) was used at 1:10. The secondary Goat anti-Mouse Alexa Fluor 594 antibody (Invitrogen Cat# A20185) was used at 1:200. Confocal images were taken using an Olympus FV1000 laser scanning confocal microscope.

## Acknowledgments

We thank Dr. Carmen Robinett, Dr. Bruce Baker, and the Developmental Studies Hybridoma Bank for Dsx antibodies, Dr. John Yoder and the Bloomington Stock Center for *Drosophila* strains, and the FlyLight Project for pBPGUw GAL4 lines. We want to thank Dr. John Yoder and Dr. Carmen Robinett for discussions of *dsx* enhancer expression, and Dr. Kohtaro Tanaka and Dr. Benjamin Vincent for comments on the manuscript.

## Competing interests

No competing interests declared

## Funding

This work was funded by NIH grants R01GM105726 and R35GM122592 to AK, by a Collaborative Supplement to NIH grant R01GM073039 to AK and MNA, and by the NSF GRFP and DDIG 1501148 to GR.

---

**Supplementary Figure 1:**
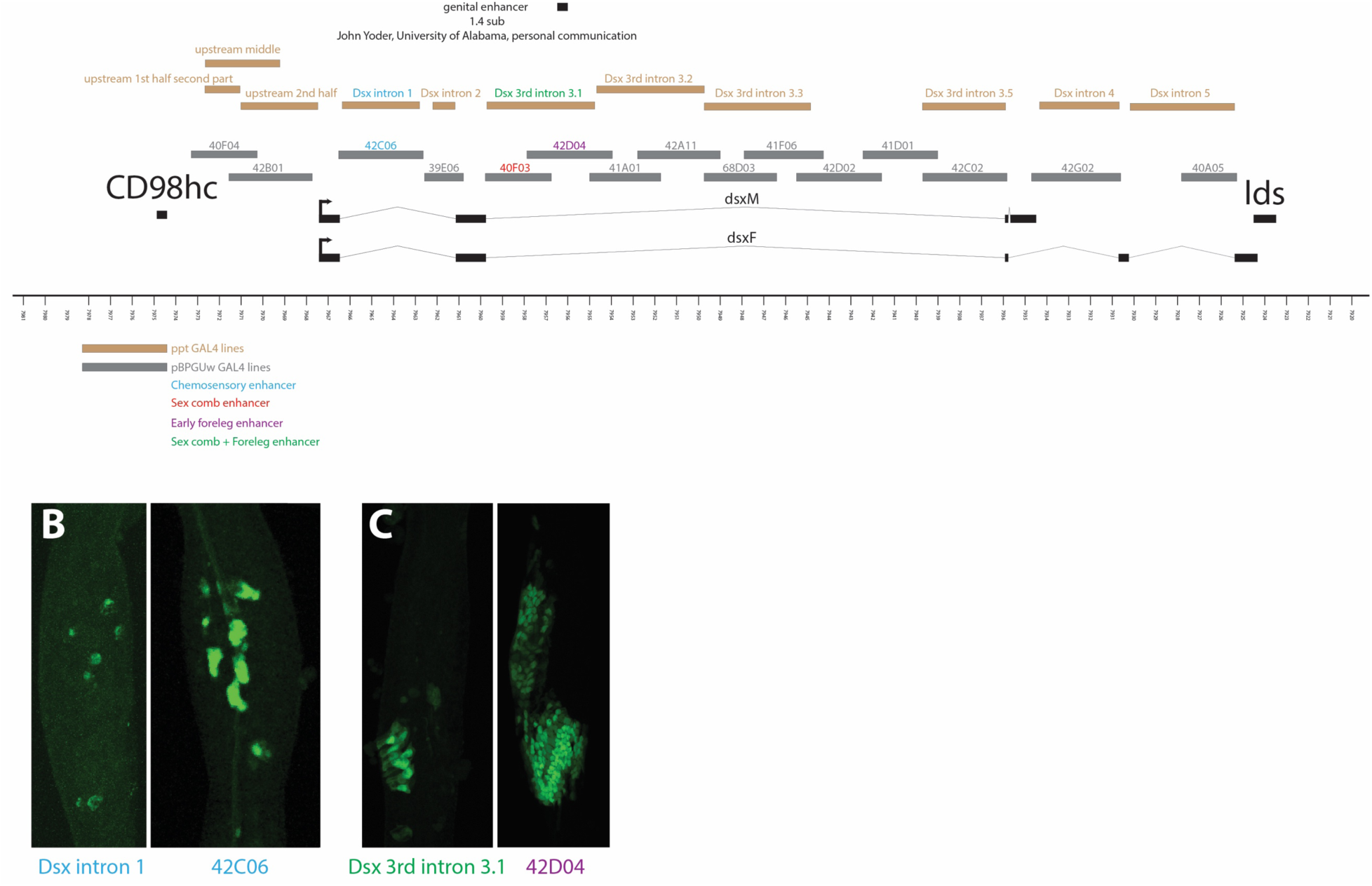
*dsx* reporter lines identify multiple regions with enhancer activity in the foreleg. To identify *dsx* foreleg enhancers, we surveyed the entire non-coding sequence of the *dsx* locus, demarcated by the flanking *CD98hc* and *lds* genes. **A**) A map of the *dsx* locus and the associated reporter lines that were analyzed. Two sets of GAL4 reporter constructs were used: a set of randomly inserted P-element lines generated in the Arbeitman lab (brown), and a set of site-specific insertions generated by (Pfeiffer et. al., 2008) (grey). *dsx* transcripts are represented by black bars (exons) and thin lines (introns). Regions that contain chemosensory, sex comb, and early foreleg enhancer are indicated in blue, red, and purple, respectively. The *dsx_intron_3.1* construct contains both the sex comb enhancer and part of the early foreleg enhancer, and is designated with green text. **B-C)** 24 hr APF male forelegs. **B**) Both the P-element integrated Dsx intron 1 (blue text, brown box) and PhiC31 integrated line 42C06 (Pfeiffer et al., 2008) (blue text, grey box) show expression in the first tarsal segment similar to the pattern of sex-specific chemosensory bristles. **C**) The P-element integrated *dsx* 3^rd^ intron 3.1 line shows expression in the sex comb region. The PhiC31 integrated line 42D04 (Pfeiffer et al., 2008) shows expression in the cells surrounding the sex comb and weak expression in the sex comb bristles.

**Supplementary Figure 2:**
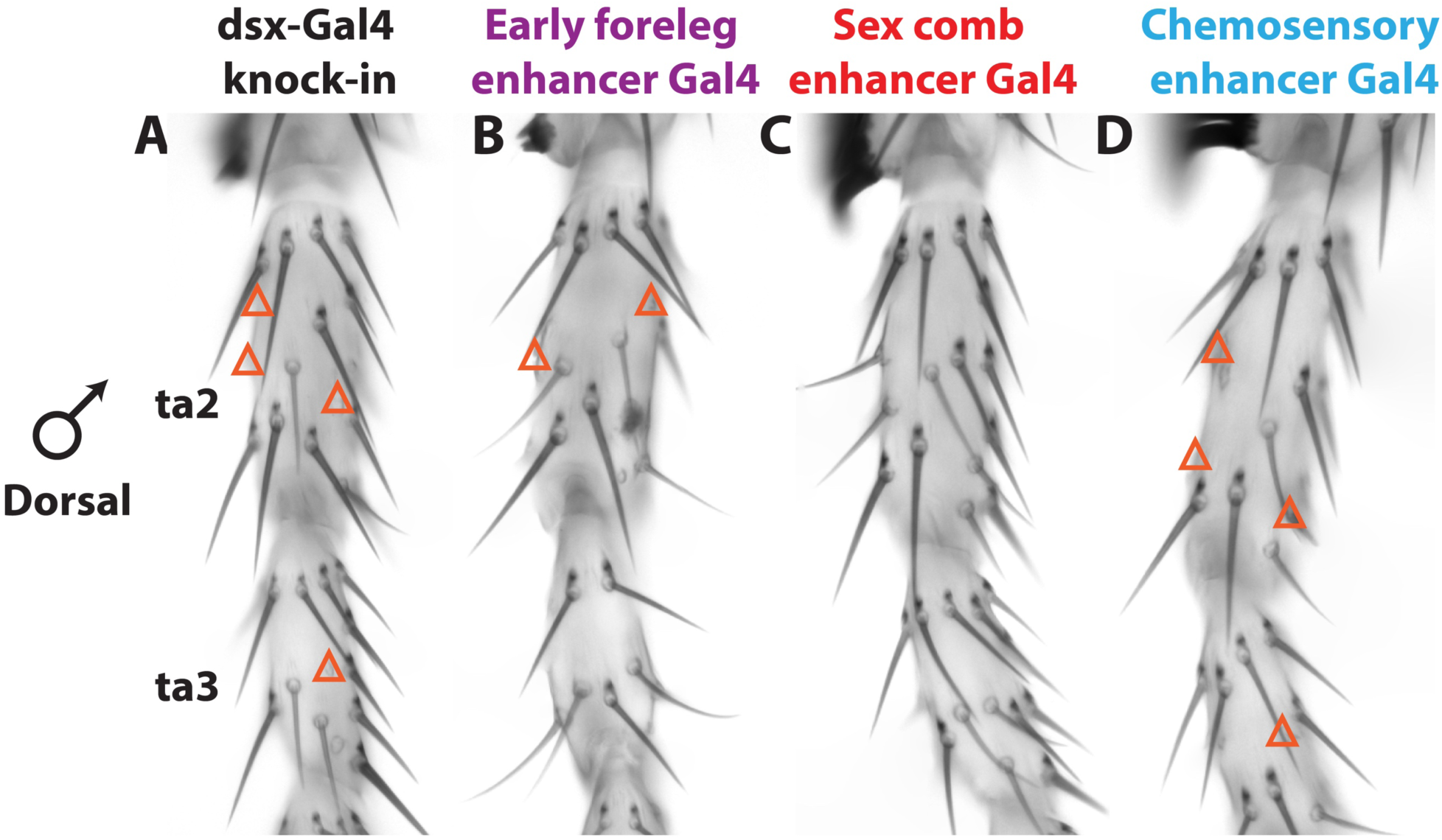
The chemosensory and early foreleg enhancers both contribute to sex-specific bristle development in ta 2 and ta3. For each enhancer, a GAL4 reporter was used to drive a UAS-*shaven* RNAi construct, truncating the bristles where that enhancer is active. Orange triangles mark chemosensory bristles that were affected in all individuals. Images of male second and third tarsal segments are shown. **A)** A GAL4 knock-in in the *dsx* gene, which likely reflects the full expression pattern of *dsx* (Robinett et al., 2010), affects three chemosensory bristles in ta2 and one chemosensory bristle in ta3. **B)** The early foreleg enhancer affects two chemosensory bristles in ta2, only one of which (left) is also affected by the *dsx*-GAL4 knock-in. **C)** The sex comb enhancer does not affect any bristles in ta2-ta3. **D)** The chemosensory enhancer affects the same chemosensory bristles in ta2-ta3 as the *dsx*-GAL4 knock-in.

**Supplementary Figure 3:**
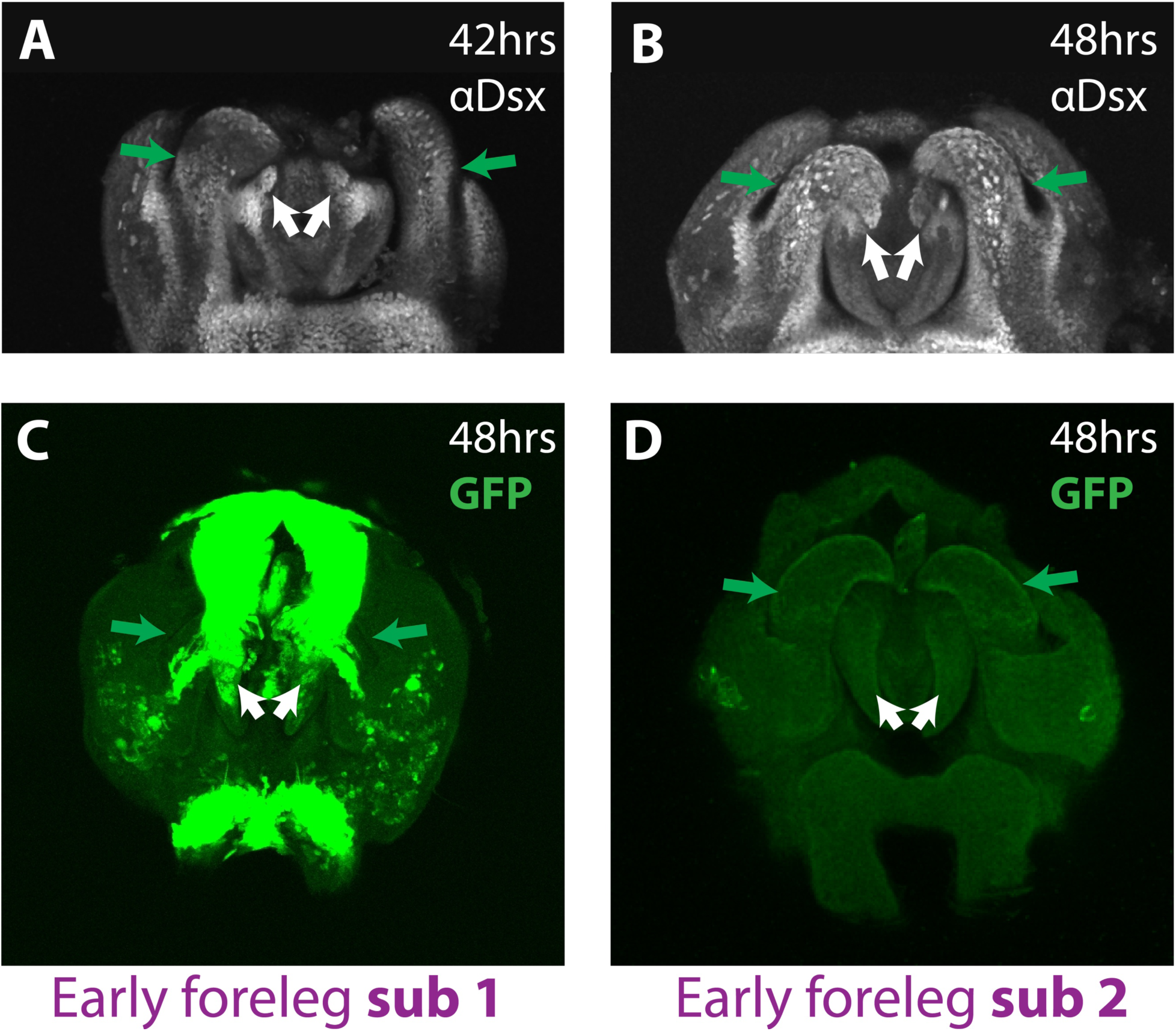
The *foreleg sub1*, but not *sub2*, enhancer fragment drives expression in male genitalia. Dsx antibody staining in pupal male genital discs at 42-48 hrs APF shows strong expression in the dorsal postgonites (green arrows) and the claspers (white arrows). The clasper expression appears to be especially prominent in bristle cells. To determine the extent of the overlap between the genital and early foreleg enhancers, we tested the ability of the *foreleg sub1* and *sub2* reporters to drive expression in the genitalia. Genital discs at 48 hrs APF carrying the *foreleg sub1* reporter showed strong expression in the dorsal postgonites (green arrows) and claspers (white arrows), similar to Dsx protein expression. No expression was seen for *foreleg sub2*.

**Supplementary Figure 4:**
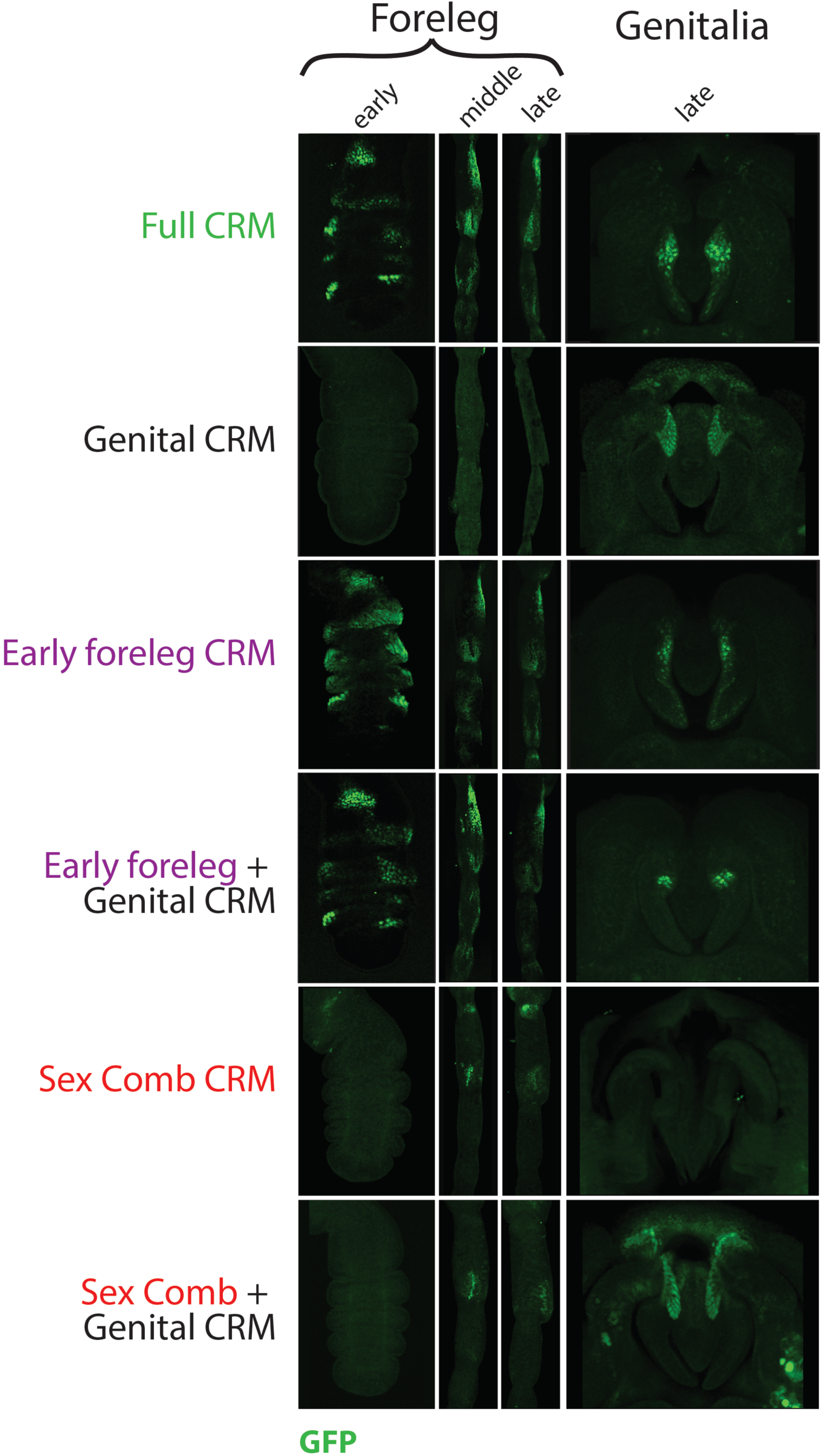
*doublesex* expression is controlled by modular enhancers. To determine whether the sex comb, genital, and early foreleg enhancers are discrete and modular, we tested the ability of non-overlapping sequences that contained each enhancer separately, as well as pairwise combinations of different enhancers, to recapitulate the expression pattern produced by all three enhancers together (“full CRM”). Early foreleg samples represent 5 hrs APF, middle 24 hrs APF, and late 43 hrs APF. The genital samples are at 48 hrs APF. GFP expression is in green. Only the constructs that contain the early foreleg enhancer show activity at 5 hrs APF. The activity of the early foreleg enhancer in the leg does not change with the addition of the genital enhancer. At 24 hrs APF, the full enhancer shows expression both in the sex comb bristle cells and in the epithelial cells surrounding the sex comb. The early foreleg enhancer shows strong expression in the epithelial cells surrounding the sex comb, but weak expression in the sex comb bristle cells. The sex comb enhancer drives expression in the sex comb bristles cells. The genital enhancer does not show any leg expression, and its inclusion does not change the spatial expression of either the sex comb or the early foreleg enhancer. At 43 hrs APF, the full enhancer shows expression in the sex comb bristle cells and the adjacent epithelial cells. The early foreleg enhancer shows strong expression in the epithelial cells surrounding the sex comb, but weak expression in the sex comb bristles. The genital enhancer does not show foreleg expression. In the genital disc at 48 hrs APF, the full enhancer is expressed in the dorsal region of the phallic sheath. The sex comb enhancer does not drive expression in the sheath, while the early foreleg enhancer drives expression in the same region as the full enhancer.

**Supplementary Table 1:**
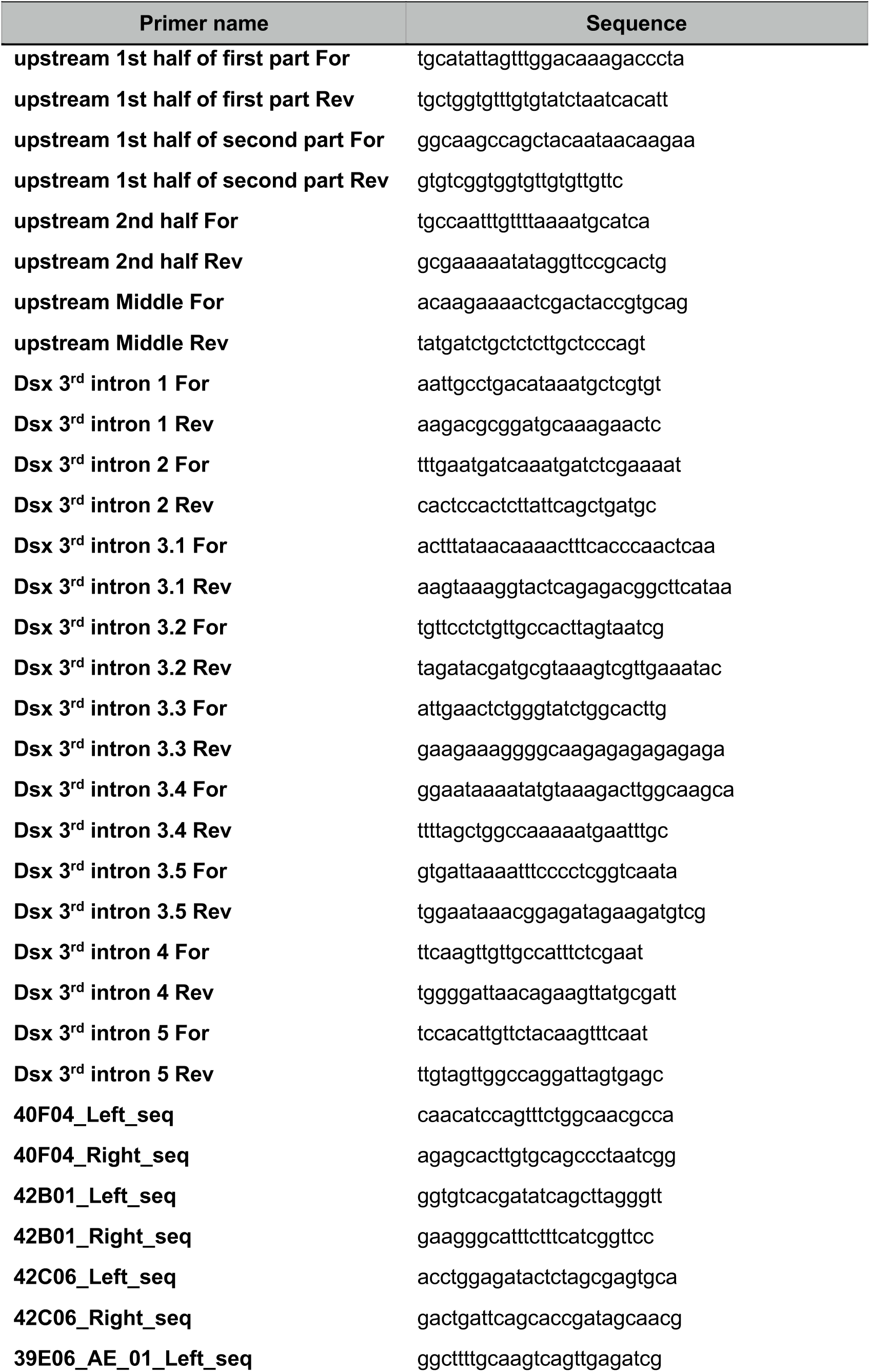
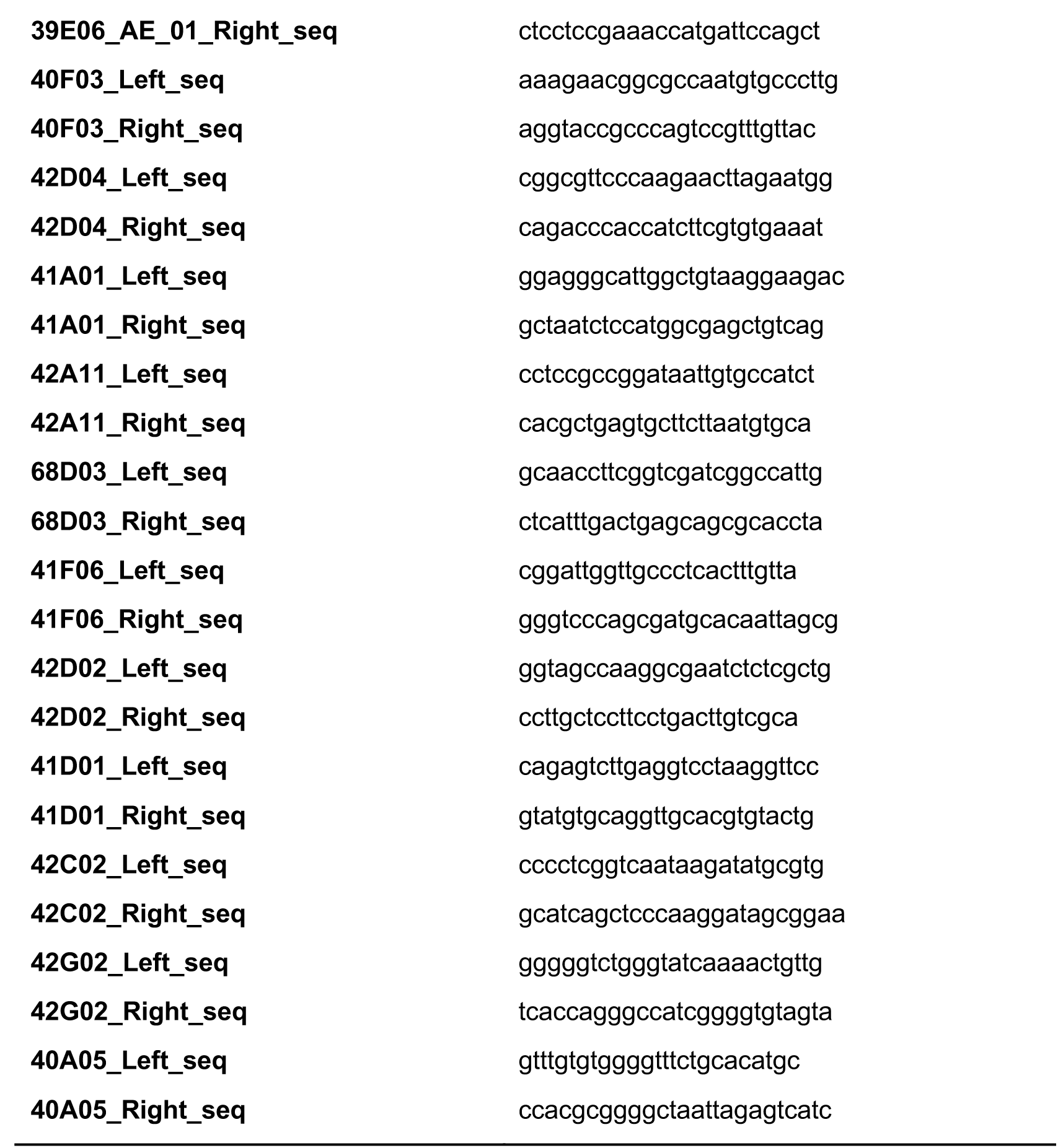
List of primers for initial enhancer screen.

**Supplementary Table 2:**
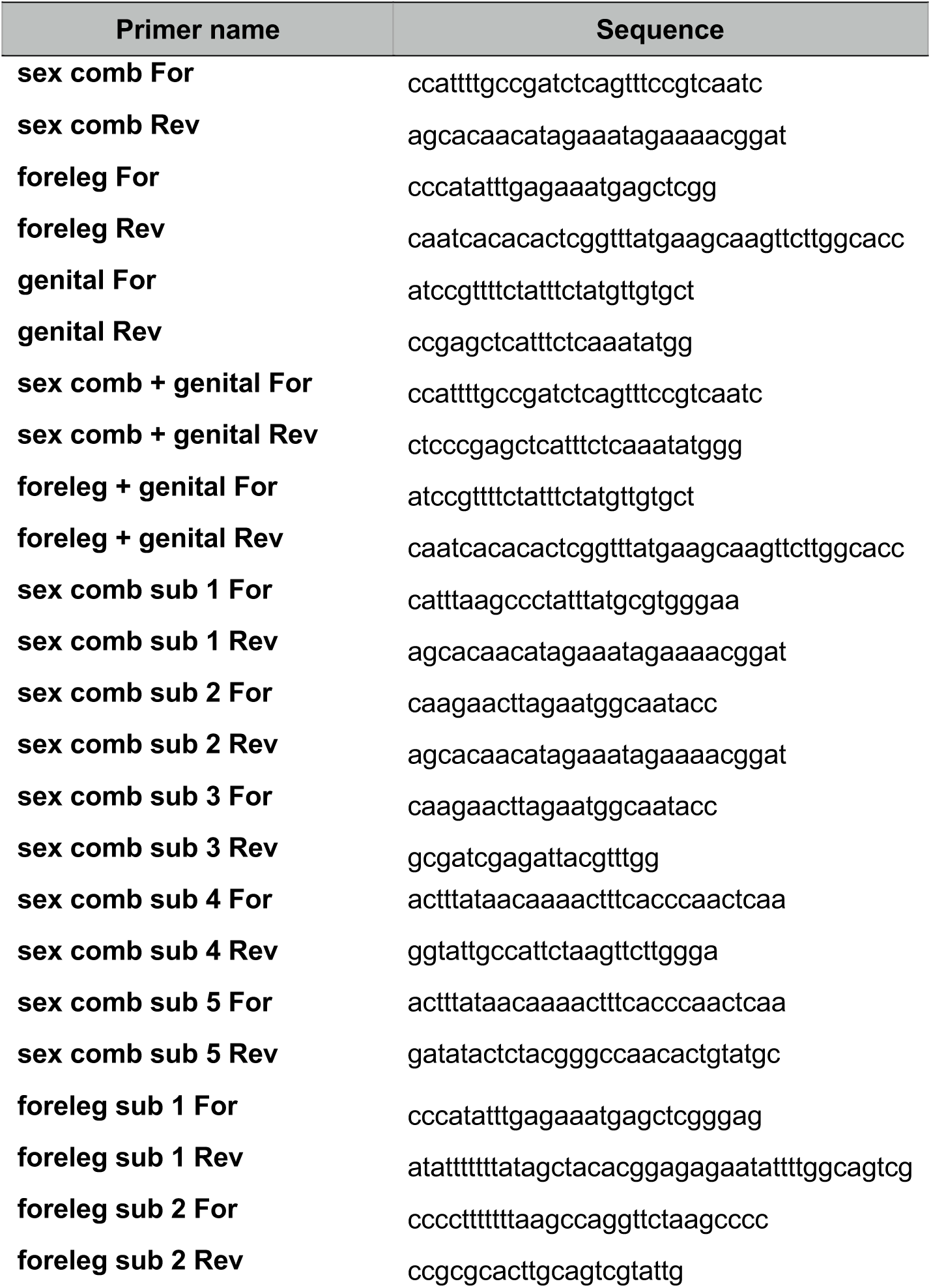
List of primers for subdivisions of sex comb and foreleg enhancers.

